# New peptides against B16F10 interfere in cell cycle of melanoma cells

**DOI:** 10.1101/2024.01.21.576573

**Authors:** Eric Almeida Xavier

## Abstract

Peptides have fantastic functions, they can act interfering in various cellular mechanisms such as the cell cycle. Furthermore, because of their high specificity, peptides can be used in antitumor therapy against specific targets. In this work we describe the *in vitro* action of four antitumor peptide against B16F10 melanoma cells, obtained from immunoglobulin genes. As a result, we show that peptides interfered in the cell cycle of B16F10 cells. In conclusion the present article describes a molecule from immunoglobulin with future potential for *in vivo* therapeutic test.

## 1. Introduction

In nature peptides have fantastic functions and capabilities, they build adaptable viral across species [1]. For example, viral pathogens that cause a group of fatal diseases or prions that are devoid of nucleic acid [2,3,4]. With the aim of exploring the antitumor capabilities of peptides as their high specificity and ability to inhibit important tumor pathways we test the potential of a molecules obtained by a group collaborator [5]. The important capacity of peptides to inhibit tumors has been previously demonstrated by our laboratory through various works [6,7,8].

The peptides were obtained from locus immunoglobulin (Ig) genes and amidated in C-terminal to increase peptide stability for potentials in vivo experiments. The selection of the sequences followed the criteria: presence of positively charged residues, net charge, isoelectric point, and alternation of hydrophobic/hydrophilic residues in the sequence, by using ExPASy Proteomics Tools Compute pI/MW and ProtParam. These selection criteria are hypothetically similar to protective molecules of natural immunity, [9]. Like, phylogenetic evolution of defensins, the natural peptides with similarity in mammalian plants, bacteria and fungi [10]. Consequently, with previous biologic action, such results justified the test against B16F10 murine melanoma tumor cells [5,6,7,8]. Finally, the peptides had in vitro antitumor activity against cell cycle in B16F10.

## 2. Material and methods

### 2.1. Cell cycle analysis with propidium iodide (PI)

Control cells were incubated with dimethylsulfoxid (DMSO) at a concentration of 20 mg/ml and treated cells were inoculated with 1 mM of peptide. Harvested tumor cells were washed twice with PBS by spinning at 300 g for 5 min and discarding the supernatant before resuspension of cells at 3 × 106 cells/mL in a cell suspension buffer (PBS + 2% fetal bovine serum, FBS; PBS + 0.1% bovine serum albumin, BSA). Cell suspensions in 500 μL buffer aliquoted in 15 mL Vbottomed polypropylene tubes received 5 ml of cold 70% ethanol dropwise with gently vortexing. Cells were fixed for at least 1 h at 4 °C prior to propidium iodide (PI) staining and flow cytometric analysis. Fixed cells were washed twice in PBS by centrifugation. One mL of PI staining solution at 50 μg/mL was added to the cell pellet. A final concentration 0.5 μg/mL in 50 μL of RNase A stock solution was also added to the cells and incubation was performed overnight (or at least 4 h) at 4 °C. Stored samples kept at 4 °C were 106 events analyzed by flow cytometry BD Accuri™ C6 Plus.

### 2.2. Cell lines and culture conditions

The murine melanoma cell line B16F10-Nex2 was originally obtained from the Ludwig Institute for Cancer Research (LICR), São Paulo branch. The cell line grew in RPMI-1640 (Gibco, Grand Island, NY) medium supplemented with 10 mM of 2-(4-(2-hydroxyethyl) piperazin-1-yl) ethane sulfonic acid (HEPES; Sigma-Aldrich, St. Louis, MO), 24 mM sodium bicarbonate, 40 mg/L gentamicin (Hipolabor, Minas Gerais, Brazil), pH 7.2, and 10% fetal bovine serum (Gibco, Grand Island, NY). Cells were cultured at 37 °C and 5% CO2 and 95% humidity in the atmosphere.

### 2.3. Selection and synthesis of peptide encoded by immunoglobulin gene

The peptides sequences used in the present work are: [7-8] with amidated C-terminal. The research exploiting the Gene database of the National Center for Biotechnology Information (NCBI, https://www.ncbi.nlm.nih.gov). According to different criteria, i.e. presence of positively charged residues, net charge, isoelectric point, and alternation of hydrophobic/hydrophilic residues in the sequence, by using ExPASy Proteomics Tools Compute pI/MW and ProtParam (http://www.expasy.org/proteomics). Selected peptides were synthesised by solid phase peptide synthesis method using a multiple peptide synthesiser (SyroII, MultiSynTech GmbH), at CRIBI Biotechnology Center (University of Padua, Italy). The purity of peptides, evaluated by analytical reverse phase HPLC, was in the 80–90% range. The peptides were solubilised in dimethyl sulfoxide (DMSO) at a concentration of 20 mg/ml and subsequently diluted in sterile distilled water for experimental use. For all experiments, controls (in the absence of peptides) contained dimethyl sulfoxide at the proper concentration.

## 3. Results and discussion

We used flow cytometry to analyze the cell cycle using propidium iodide DNA staining. We observed a difference between peptides named ER1, L12P, W12K and G10S compared whit control PBS+DMSO treated cells. Where was verified an interfere in cell cycle (Figure. 3 A and B). Lastly, the effect on cell cycle could be explained by the interaction of peptides with cycle molecules, thereby the tumor cells are unable to precisely make the check point between the G1 and S phases and as a consequence of this, there was an expansion of the S phase and a shrinkage of G2/M phases observed in (Figure. 1 A-C) [11,12]; in the future this result can be elucidated by Western blotting of the molecules involved in the cell cycle.

**Figure 1.**
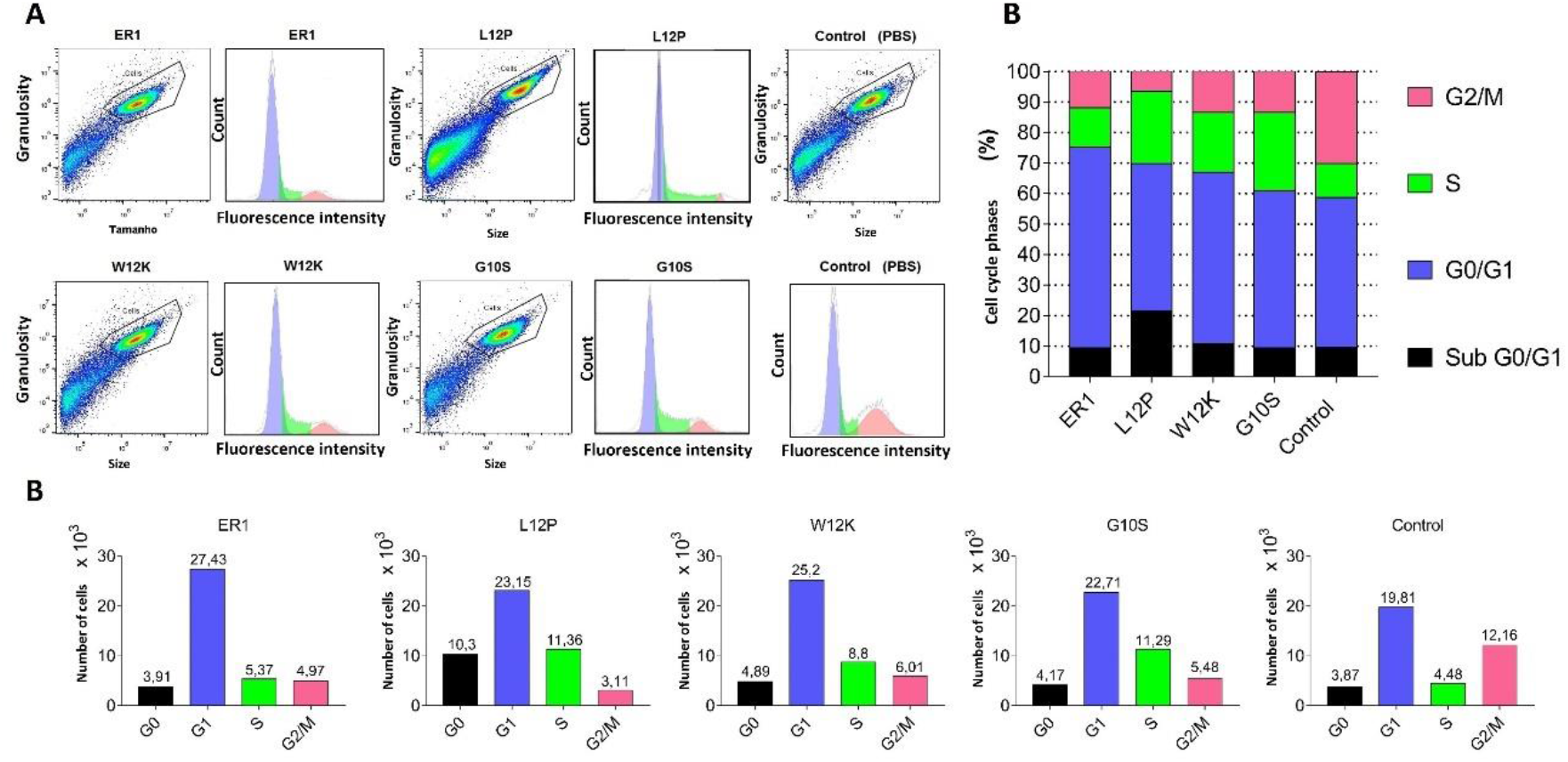
(**A**) 10^6^ B16F10 cells were treated with ER1, L12P, W12K and G10S after analyzed by flow cytometer at incubation with propidium iodide (PI) solution. (**B**) Percentage of each phase of the cycle in G1, S and G2 phases. (**C**) PI staining shows different intensities in the population of treated cells with difference in cell cycle pattern in comparison to PBS + DMSO control. The graphs and analyzes were made using FlowJo software.

## 4. Conclusion

Finally, this study will benefit the society to obtaining peptides sequences that have therapeutic action against melanoma and possibly for many other types of cancer. Thus, we homologate the peptide for future in vivo assays. To acts on cell viability, by inhibition of cell cycle. And because that causing a delay in tumor progression.

## Compliance with ethical standards

## Acknowledgments

I would like to thank my Professor Maurício da Silva Baptista for his confidence in my work. I would also like to thank the São Paulo Government Research Support Foundation (FAPESP) for funding.

## Consent for Publication

We authorize the full disclosure of the manuscript text and data.

## References

[1] Gonçalves D, Prado RQ, Xavier EA, de Oliveira NC, Guedes PMDM, da Silva JS, et al. Corrigendum to Immunocompetent Mice Model for Dengue Virus Infection. Scientific World Journal. 2018; 5268929.

[2] Xavier, Eric Almeida. Prions: the danger of biochemical weapons. Food Science and Technology. 2014, 34(3): 433–440.

[3] Eric Almeida Xavier. Prion/virus the danger of biological weapons. World Journal of Biology Pharmacy and Health Sciences. 2022, 10(03): 031–035.

[4] Eric Almeida Xavier. Prions and virus pathogen gain-of-function. World Journal of Advanced Research and Reviews. 2022, 15(03): 169–175.

[5] Eric Almeida Xavier and Fabrício Castro Machado. L18R a peptide with potential against melanoma. Open Access Research Journal of Biology and Pharmacy. 2022, 04(02): 022–027.

[6] Eric Almeida Xavier. G10S an immunoglobulin peptide against melanoma. World Journal of Advanced Research and Reviews. 2022, 14(01):385–390

[7] Eric Almeida Xavier. An in vitro test of new peptide against melanoma. World Journal of Advanced Research and Reviews. 2022, 14(01): 024–028.

[8] Eric A Xavier, Fabricio C Machado, Luiz Rodolpho RG Travassos. Immunoglobulin peptide against melanoma. World Journal of Advanced Pharmaceutical and Life Sciences, 2022, 02(01): 009–013.

[9] Litman GW, Cannon JP, Dishaw LJ. Reconstructing immune phylogeny: new perspectives. Nat Rev Immunol. 2005; 5(11): 866–79.

[10] Carvalho AeO, Gomes VM. Plant defensins and defensin-like peptides - biological activities and biotechnological applications. Curr Pharm Des. 2011; 17(38): 4270–93.

[11] Baldin V, Lukas J, Marcote MJ, Pagano M, Draetta G. Cyclin D1 is a nuclear protein required for cell cycle progression in G1. Genes Dev. 1993; 7(5): 812–21.

[12] Suski JM, Braun M, Strmiska V, Sicinski P. Targeting cell-cycle machinery in cancer. Cancer Cell. 2021; 39 (6): 759–78.

